# Clinical factors associated with SFTS diagnosis and severity in cats

**DOI:** 10.1101/2024.03.06.583784

**Authors:** Hiromu Osako, Qiang Xu, Takeshi Nabeshima, Jean Claude Balingit, Khine Mya Nwe, Fuxun Yu, Shingo Inoue, Daisuke Hayasaka, Mya Myat Ngwe Tun, Kouichi Morita, Yuki Takamatsu

**Author notes:** **Corresponding author contact information** Yuki Takamatsu; Department of Virology, ITM-NU, Sakamoto1-12-4 Nagasaki, 852-8523, Japan. These authors contributed equally (Hiromu Osako, Qiang Xu).

## Abstract

Severe fever with thrombocytopenia syndrome (SFTS) is a potentially fatal tick-borne zoonosis caused by SFTS virus (SFTSV). In addition to tick bites, animal-to-human transmission of SFTSV has been reported, but little is known about feline SFTSV infection. In this study, we analyzed data on 187 cats with suspected SFTS to identify biomarkers for SFTS diagnosis and clinical outcome., Body weight, red and white blood cell and platelet counts, serum aspartate aminotransferase and total bilirubin levels, were useful for SFTS diagnosis, and alanine aminotransferase, aspartate aminotransferase and serum SFTSV RNA levels were associated with the clinical outcome. We developed a scoring model to predict SFTSV infection. In addition, we performed a phylogenetic analysis to reveal the relationship between disease severity and viral strain. This study provides comprehensive information on feline SFTS and could contribute to the protection of cat owners, community members, and veterinarians from the risk of cat-transmitted SFTSV infection.

## Introduction

Severe fever with thrombocytopenia syndrome (SFTS) is a viral infection first discovered in China in 2010.^1^ It has subsequently been identified in other countries in East Asia, including South Korea^2^ and Japan,^3^ and it has recently emerged in South East Asia including Vietnam,^4^ Taiwan,^5^ Pakistan,^6^ and Myanmar.^7^ As the endemic region has expanded, SFTS has become an important public health concern.

SFTS is caused by SFTS virus (SFTSV), a member of the genus *Bandavirus* in the Phenuiviridae family.^1,8^ The main route of SFTSV transmission is tick bites, with some cases transmitted by exposure to bodily fluids from infected individuals.^9,10^ Recently, cats have shown similar clinical manifestations to human cases,^11^ and transmission from cats to humans has been reported.^12–14^ Moreover, a case-control study of human SFTS in China showed an association between cat ownership and risk of infection.^15^ Therefore, cats might be an important source of SFTSV transmission to humans.

Regarding human SFTS cases, the risk of infection is high in adults aged over 50 years, whereas there is no difference according to sex.^16^ In the early stages of SFTS, patients present with flu-like symptoms such as fever, fatigue, headache, myalgia, and gastrointestinal symptoms (loss of appetite, nausea, vomiting, and diarrhea).^17,18^ The disease progresses to lymphadenopathy, hemorrhagic signs, and involvement of the central nervous system is observed approximately 5 days after onset and persists for 1 to 2 weeks.^17,18^ During this phase, most fatal cases develop multiple organ failure (MOF).^17^ Age, sex, and elevated levels of serum aspartate aminotransferase (AST), lactate dehydrogenase (LDH), creatine kinase (CK), and creatine kinase MB isoenzyme (CK-MB) are associated with the risk of death in SFTS.^17^ Although the case fatality rate of SFTS has been reported to be up to 30%,^19^ no effective treatment has yet been developed.^20^ Therefore, it is imperative to understand the pathogenicity of SFTS and develop countermeasures.

The incidence of feline SFTS in Nagasaki is among the highest in Japan.^21^ Notably, Nagasaki is an area with numerous communally owned cats (cared for by members of the community with no specific owner) live, indicating that cats are a potential risk factor for SFTSV transmission to the surrounding community. Cat-to-human transmission has also been reported in Nagasaki in recent years (unpublished data). However, currently no detailed information is available on the clinical manifestations and laboratory parameters for SFTS diagnosis or prognostic factors in cats. Veterinarians and cat owners sometimes have difficulty in confirming the diagnosis in cases of suspected SFTS in cats, and our laboratory has received numerous requests for laboratory confirmation of the diagnosis. Therefore, in this study, we aimed to identify biomarkers for SFTS diagnosis and clinical outcomes using samples from cats with suspected SFTS provided by several veterinary hospitals in Nagasaki Prefecture. This study identified biomarkers for SFTSV infection, contributing to the protection of owners, surrounding persons, veterinarians, and healthcare professionals from the risk of SFTSV infection in cats.

## Materials and methods

### Cat specimens

We obtained serum and swab specimens from 221 cats with suspected SFTS from animal hospitals in Nagasaki Prefecture between March 2018 and January 2024. Among them, the necessary information was not available in 24 cases; therefore the analysis included 187 cases that were sampled within 7 days from onset and were identified as SFTSV-positive or negative. The specimens including sera, oral swabs, anal swabs, and eye swabs, were stored and transported at 4°C to the department of Virology, Institute of Tropical Medicine, Nagasaki University. Data sheets, including information on sex, age, weight, clinical signs, blood test results (hematology and biochemistry), and sampling region, were provided together with the specimens.

### SFTSV RNA detection

The procedure for SFTSV genome detection in cat specimens using real-time quantitative polymerase chain reaction (RT-qPCR) has been described.^22,23^ Briefly, RNA was extracted using Isogen-LS (Nippon Gene) and RT-qPCR was performed using the One-Step PrimeScript RT-PCR Kit (Takara Bio) on a 7500 Real-Time RT-PCR System (Applied Biosystems). Specific primers and probes (Table S1) were developed based on the consensus genome sequence of the RNA-dependent RNA polymerase (RdRp) -coding region in the SFTSV-L segment.^22^ The viral genome copy number was calculated based on the standard control ranged from 10^2^ to 10^8^ genome copies/5-μL.

### Cells and viruses

Vero E6 (African green monkey kidney) cells were cultured in Dulbecco’s modified Eagle’s medium (DMEM; Fujifilm Wako Pure Chemical) supplemented with 10% fetal bovine serum (FBS; Biowest), P/S; 100 IU/mL penicillin and 100 μg/mL streptomycin (PS; Sigma-Aldrich) at 37°C in a humidified atmosphere containing 5% CO_2_. In this study, the SFTSV YG-1 strain, which was isolated from serum specimens of a Japanese patient with SFTS,^3^ was used to prepare antigens to detect cat SFTSV-reacting immunoglobulin G (IgG).

The SFTSV viral titers were assessed using a focus-forming assay as described previously,^22,23^ with minor adjustments. A monolayer of Vero E6 cells cultured in 96-well plates were exposed to a ten-fold serially diluted SFTSV with DMEM and were incubated at 37°C for 1 hour. Then, an overlay medium, DMEM supplemented with 2% FBS and 1.25% methylcellulose 4000 (Fujifilm Wako Pure Chemical), was added for 5 days and fixed in a 4% Paraformaldehyde Fix Solution (Fujifilm Wako Pure Chemical). Viral foci were identified using monoclonal antibodies (mAb: 4A10) targeting the SFTSV-N protein,^24^ and then treated with goat anti-mouse IgG conjugated with horseradish peroxidase (American Qualex International), followed by the addition of the substrate: a 3,3′-diaminobenzidine (DAB) tablet (Fujifilm Wako) dissolved in phosphate-buffered saline (PBS) and 30% H_2_O_2_ (Sigma-Aldrich), according to the manufacturer’s instructions. Viral titers were determined as focus-forming units per milliliter (ffu/mL). All experiments involving infectious SFTSV were conducted in a Biosafety Level 3 facility at the Institute of Tropical Medical at Nagasaki University, following Nagasaki University guidelines and based on the national laws.

### Virus isolation

Virus isolation was performed using SFTSV RNA-positive serum specimens which RNA copy number>10^5^/5μL (27 specimens from 88 cases), collected during January 2021 to December 2023, as previously described.^3,23,25^ Due to the limitation of storage condition, samples from March 2018 to December 2020 were not available for virus isolation. Briefly, T25 flasks of Vero E6 cells were inoculated with 50 μL of serum specimens with DMEM devoid of FBS, and incubated at 37°C for 1 hour. Subsequently, DMEM with 2% FBS and P/S were added, and the cells were then incubated at 37°C in 5% CO_2_ for 7 days. Supernatants were collected and 500 μL of them was inoculated into fresh Vero E6 cells in T75 flasks as 2nd passages.

### Genome sequence

Viral RNA was extracted from infectious culture fluids for RT-qPCR to confirm virus isolation. The complementary DNA (cDNA) synthesis was achieved by using ReverTra Ace-α-(Toyobo) with random primer (Takara Bio). Subsequently, the M segment genes were amplified using KOD One PCR Master Mix (Toyobo) with specific primer pairs (Table S1). The amplicons-containing were excised after gel-electrophoresis and extracted DNA by using Wizard SV Gel and PCR Clean-Up System (Promega). The amplicon DNAs were further analyzed for Sanger sequencing with the QuantumDye Terminator Cycle Sequencing Kit (Tomy Digital Biology) and specific primers (Table S1) upon ABI 3500 Genetic Analyzer (Applied Biosystems). The genome sequence of the M segment-coding region has been deposited in GenBank (accession numbers: PP273701–PP273716).

### Phylogenetic analysis

All SFTSV M-segment RNA sequences retrieved from the US National Center for Biotechnology Information (NCBI) database were used for phylogenetic analysis and clustered using CD-HIT-EST (version 4.8.1).^26^ Subsequently, the sequences obtained in this study and in our previous studies^23^ were combined with the clustered dataset, sequence alignment was conducted using MAFFT (version 7.520), a multiple sequence alignment program,^27^ and phylogenetic trees were generated for the M segment genes using the maximum likelihood (ML) method based on the bootstrap approach with 1000 replications in MEGA 11.^28^

### SFTSV antibody detection

Cat serum specimens with negative RT-qPCR results from 34 cases collected during January 2021 to December 2023 were analyzed for SFTSV- reacting IgG, as previously described.^29,30^ Due to the limitation of storage condition, samples from March 2018 to December 2020 were not available for antibody detection. Briefly, Vero E6 cells were infected with the YG1 strain (MOI = 0.1) for 5 days, harvested by trypsinization, washed with PBS, and blended with uninfected Vero E6 cells at a 1:3 ratio. The cell mixture was spotted onto 12-well glass slides (Matsunami Glass Ind.), air-dried under UV irradiation for 1 hour, and fixed with pre-cooled acetone. These immunofluorescence assay (IFA) antigen slides were stored at −30°C until use. The slides were thawed to room temperature and dried before use. The cat serum specimens were heat-inactivated at 56°C for 30 minutes. Then the serum samples were serially diluted 2-fold with PBS from 1:10 to 1:10240 and 20 μL aliquots were applied to the slides. The mAb 4A10 against the SFTSV-N protein was serially diluted 2-fold with PBS from 1:100 to 1:102400 as a positive control, and PBS was used as a negative control. The slides were incubated in a humid environment at 37°C for 1 hour. They were then washed three times with PBS, and the slides were reacted with FITC-conjugated goat anti-feline IgG (H + L) antibody (Thermo Fisher Scientific) diluted with PBS at a ratio of 1:500 for serum specimens and the negative control, and Alexa Fluor 594 conjugated donkey anti-mouse IgG (H + L) antibody (Abcam) as a positive control. Simultaneously, the nuclei were counterstained with Hoechst stain at a ratio of 1:2000. The slides were then incubated in a humid environment at 37°C for 1 hour. After washing three times with PBS, glass covers were applied and the slides were examined under a fluorescence microscope (Keyence BZ-X810).

### Statistical analysis

Data analysis and visualization were performed using R 4.2.2.^31^ The Wilcoxon rank-sum test was used to compare continuous variables, such as individual data, clinical manifestations, laboratory parameters, and viral RNA loads, between groups. Fisher’s exact test was used to compare categorical variables, such as sex, clinical signs, and clinical outcome, between groups. Pearson correlation coefficients were used to assess correlations between continuous variables. In the correlation matrix analysis, Spearman correlation coefficients between variables were calculated with false-discovery rate (FDR) correction and were visualized using the “psych” and “corrplot” functions in R (The R Foundation for Statistical Computing, Vienna, Austria). The Hosmer– Lemeshow test was used to evaluate a regression model. A scoring model was constructed as described previously.^32^ Missing values were excluded from the analysis. P-values < 0.05 were considered statistically significant.

## Results

### Comparison of SFTSV-positive and SFTSV-negative cases

The basic characteristics of cats with suspected SFTS are shown in Table 1. There were no differences in sex (p = 0.646), history of tick bites (p = 0.130), or rearing environment (p = 0.606) between SFTSV-positive and SFTSV-negative cases. The proportion with diarrhea (p = 1.000) and lethargy (p=0.344) did not significantly differ between SFTSV-positive and SFTSV-negative cases. Although there was no significant difference in vomiting (p=0.051) and outcome (p = 0.063) between SFTSV-positive and SFTSV-negative cases, vomiting was more frequently observed in SFTSV-positive cases (22.1%) than in SFTSV-negative cases (10.0%), and the mortality in the SFTSV-positive group (46.8%) was higher than that in the SFTSV-negative group (20.0%).

**Table 1.**
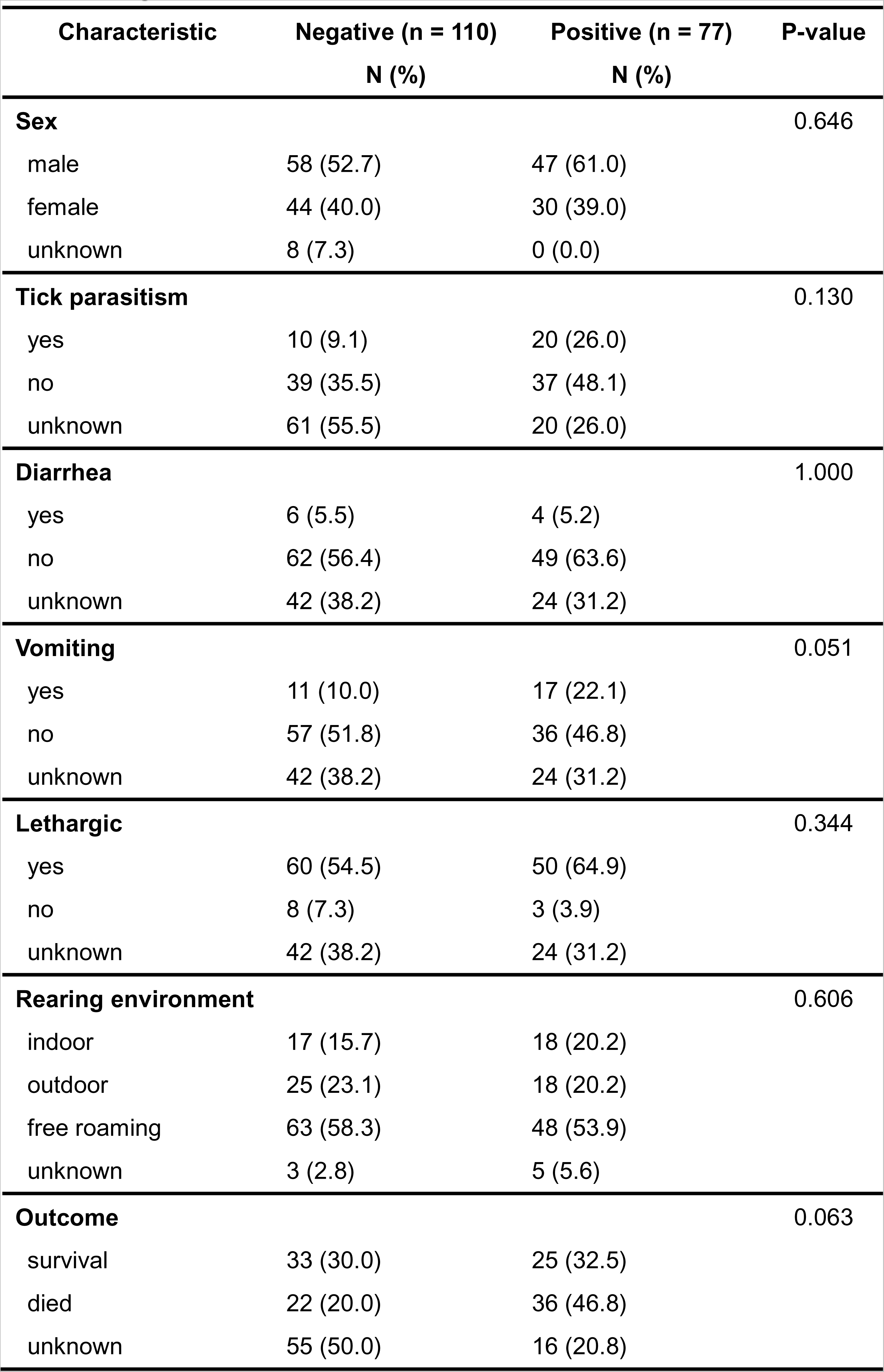
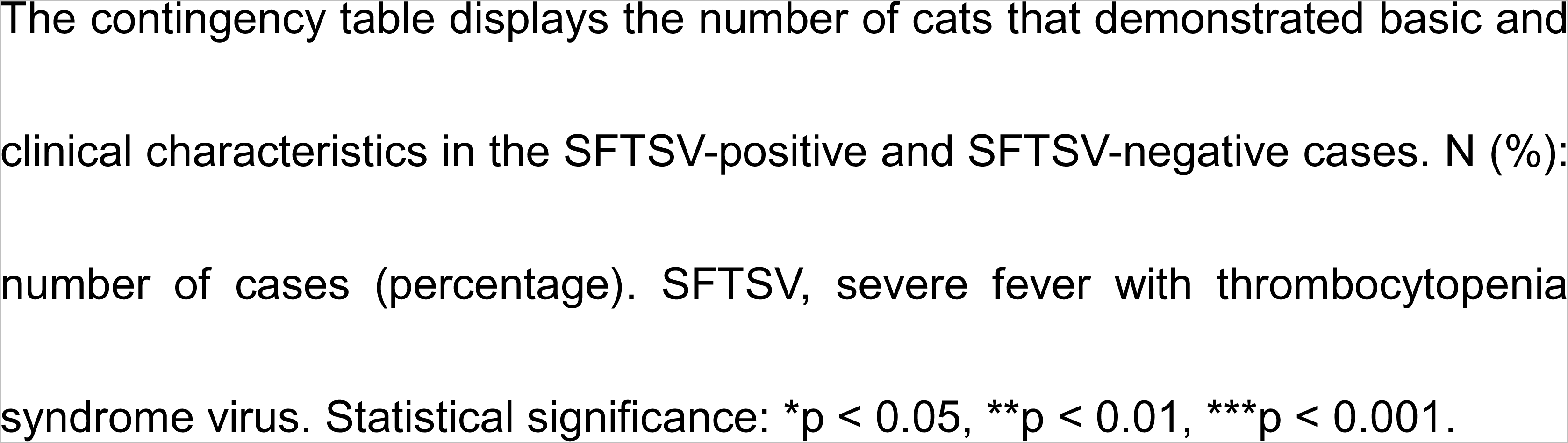
Differences in clinical parameters between SFTSV-positive and SFTSV-negative cases.

| Characteristic | Negative (n = 110)<br>N (%) | Positive (n = 77)<br>N (%) | P-value |
| --- | --- | --- | --- |
| **Sex** |  |  | 0.646 |
| male | 58 (52.7) | 47 (61.0) |  |
| female | 44 (40.0) | 30 (39.0) |  |
| unknown | 8 (7.3) | 0 (0.0) |  |
| **Tick parasitism** |  |  | 0.130 |
| yes | 10 (9.1) | 20 (26.0) |  |
| no | 39 (35.5) | 37 (48.1) |  |
| unknown | 61 (55.5) | 20 (26.0) |  |
| **Diarrhea** |  |  | 1.000 |
| yes | 6 (5.5) | 4 (5.2) |  |
| no | 62 (56.4) | 49 (63.6) |  |
| unknown | 42 (38.2) | 24 (31.2) |  |
| **Vomiting** |  |  | 0.051 |
| yes | 11 (10.0) | 17 (22.1) |  |
| no | 57 (51.8) | 36 (46.8) |  |
| unknown | 42 (38.2) | 24 (31.2) |  |
| **Lethargic** |  |  | 0.344 |
| yes | 60 (54.5) | 50 (64.9) |  |
| no | 8 (7.3) | 3 (3.9) |  |
| unknown | 42 (38.2) | 24 (31.2) |  |
| **Rearing environment** |  |  | 0.606 |
| indoor | 17 (15.7) | 18 (20.2) |  |
| outdoor | 25 (23.1) | 18 (20.2) |  |
| free roaming | 63 (58.3) | 48 (53.9) |  |
| unknown | 3 (2.8) | 5 (5.6) |  |
| **Outcome** |  |  | 0.063 |
| survival | 33 (30.0) | 25 (32.5) |  |
| died | 22 (20.0) | 36 (46.8) |  |
| unknown | 55 (50.0) | 16 (20.8) |  |

The contingency table displays the number of cats that demonstrated basic and clinical characteristics in the SFTSV-positive and SFTSV-negative cases. N (%): number of cases (percentage). SFTSV, severe fever with thrombocytopenia syndrome virus. Statistical significance: \* $p < 0.05$ , \*\* $p < 0.01$ , \*\*\* $p < 0.001$ .

The differences in laboratory parameters between SFTSV-positive and SFTSV-negative groups are summarized in Figure 1 (and Table S2). Age and body temperature did not differ significantly between the SFTSV-positive and SFTSV-negative groups. However, body weight was significantly higher in the SFTSV-positive group than SFTSV-negative group (p = 0.046). The SFTSV- positive group had a significantly higher median red blood cell (RBC) count (p = 0.004) and significantly lower white blood cell (WBC) (p < 0.001) and platelet (PLT) (p = 0.014) counts. Additionally, aspartate aminotransferase (AST) and total bilirubin (TBil) levels of SFTSV-positive group were significantly higher than those of SFTSV-negative group (AST: p = 0.034, TBil: p = 0.005).

**Figure 1.**
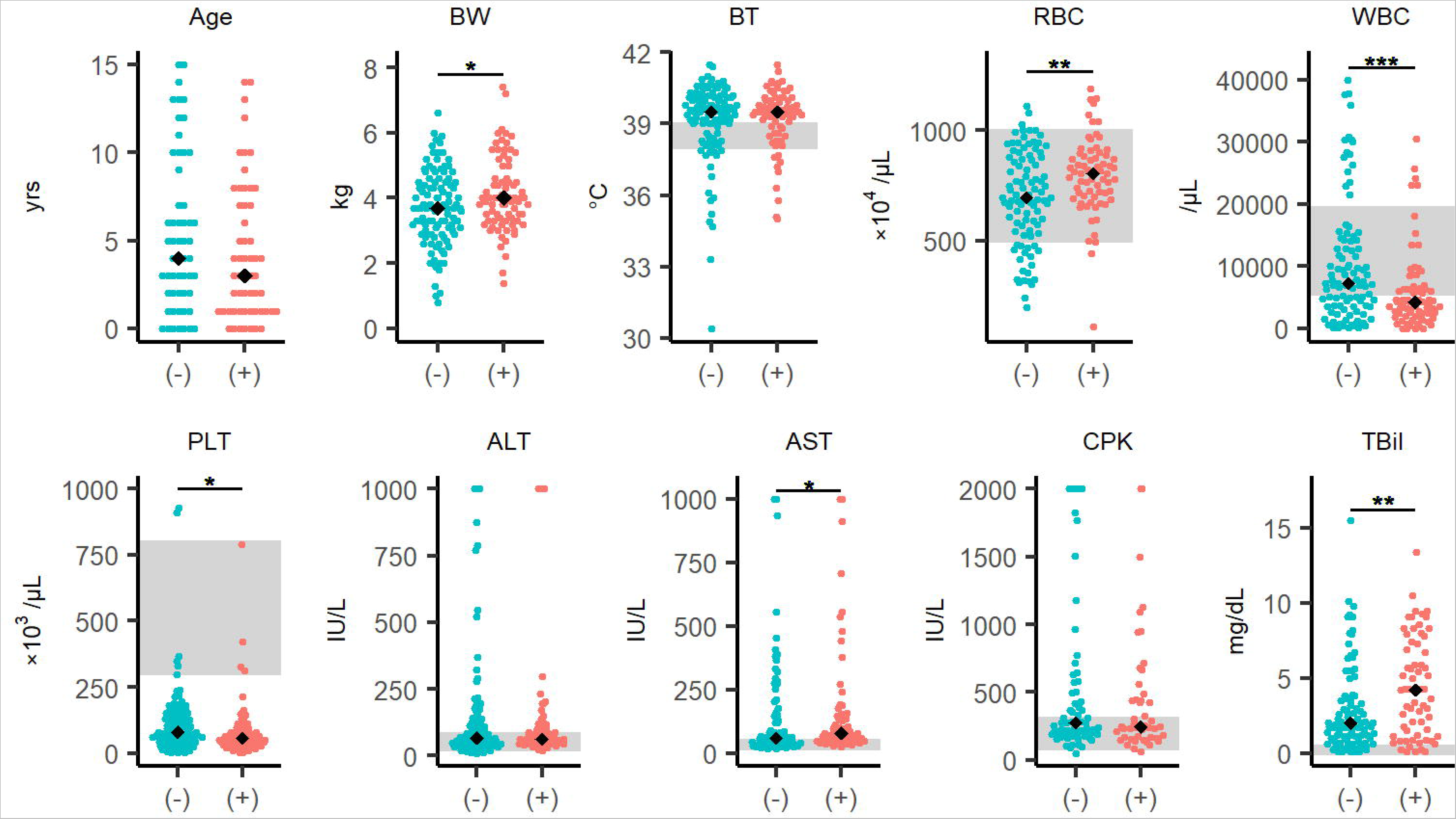
Differences in clinical characteristics between SFTSV-positive and SFTSV-negative cases. Comparison of individual data and laboratory parameters between the SFTSV- negative and SFTSV-positive groups. The analysis included 187 cases that were sampled within 7 days from onset and were identified as SFTSV-positive or negative. The black diamonds represent the median values in each group. The blue circles represent the SFTSV-negative group, and the red circles represent the SFTSV-positive group. The gray areas represent the reference ranges for body temperature, RBC, WBC, PLT, ALT, AST, CPK, and TBil. “(−)” and “(+)” stand by “SFTSV-negative” and “SFTSV-positive,” respectively. Each variable was compared using the Wilcoxon rank-sum test. The statistical significance is shown: *p < 0.05, **p < 0.01, and ***p < 0.001. See also Table S2. Abbreviations: ALT, alanine aminotransferase; AST, aspartate aminotransferase; BT, body temperature; BW, body weight; CPK, creatine phosphokinase; PLT, platelets; RBC, red blood cells; TBil, total bilirubin; WBC, white blood cell(s).

### Differences in epidemiological and laboratory parameters in the surviving and dead cases

The epidemiological differences between surviving and dead cases are summarized in Table 2. There were no statistically significant differences in sex, tick bite status, diarrhea, vomiting, lethargy, or rearing environment between the surviving and dead cases.

**Table 2.**
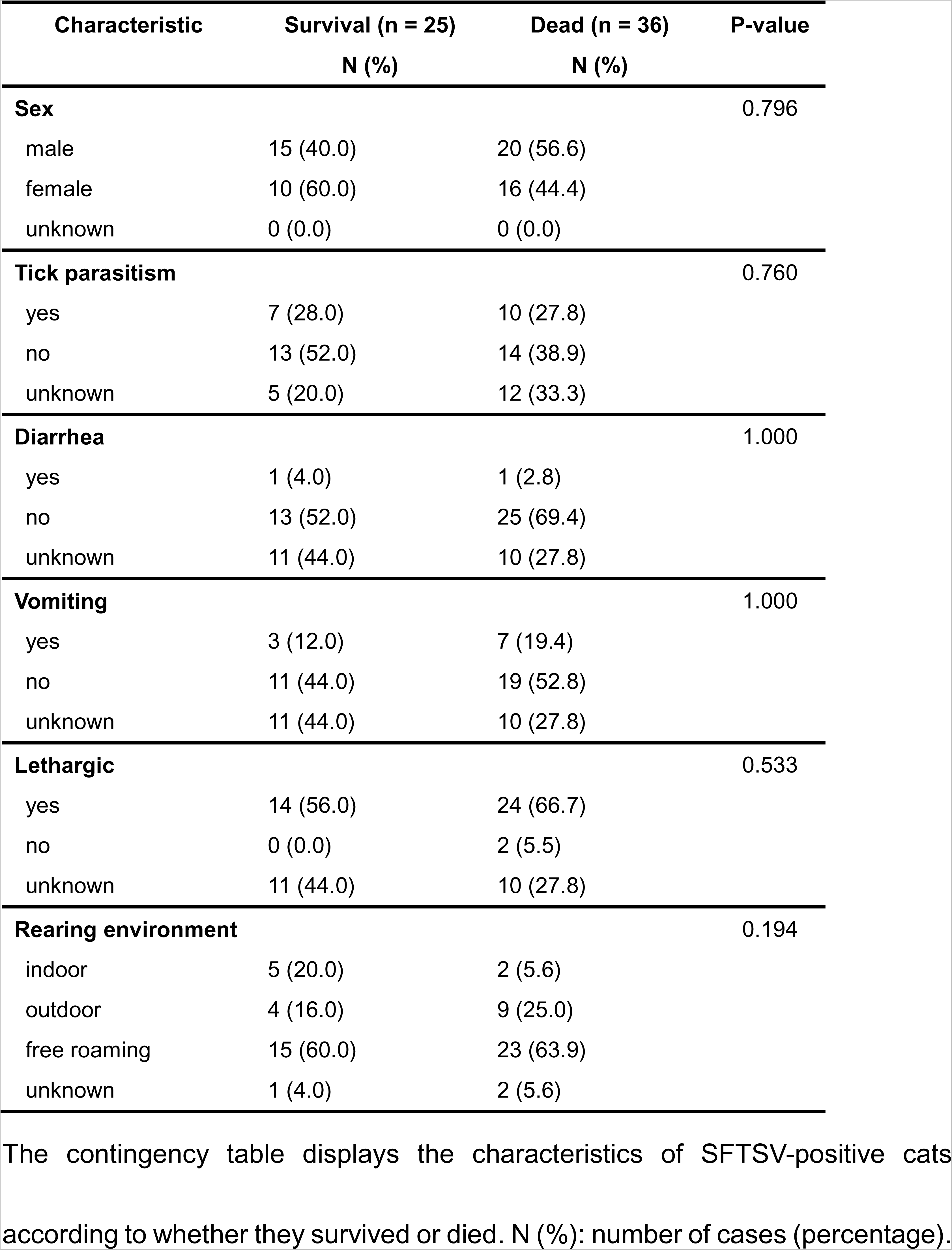
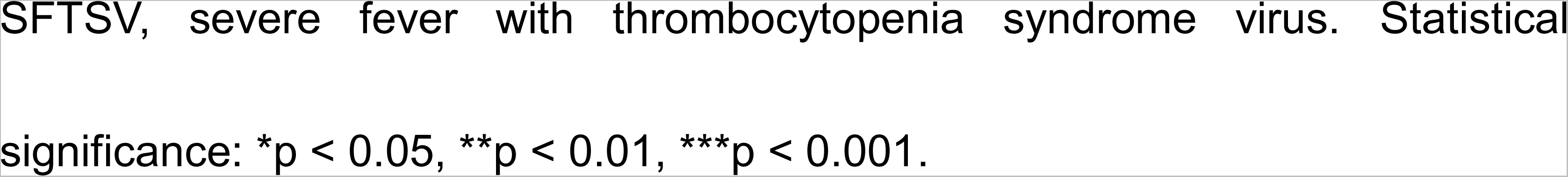
Differences in clinical parameters between survival and dead in SFTSV-positive cases.

| Characteristic | Survival (n = 25)<br>N (%) | Dead (n = 36)<br>N (%) | P-value |
| --- | --- | --- | --- |
| **Sex** |  |  | 0.796 |
| male | 15 (40.0) | 20 (56.6) |  |
| female | 10 (60.0) | 16 (44.4) |  |
| unknown | 0 (0.0) | 0 (0.0) |  |
| **Tick parasitism** |  |  | 0.760 |
| yes | 7 (28.0) | 10 (27.8) |  |
| no | 13 (52.0) | 14 (38.9) |  |
| unknown | 5 (20.0) | 12 (33.3) |  |
| **Diarrhea** |  |  | 1.000 |
| yes | 1 (4.0) | 1 (2.8) |  |
| no | 13 (52.0) | 25 (69.4) |  |
| unknown | 11 (44.0) | 10 (27.8) |  |
| **Vomiting** |  |  | 1.000 |
| yes | 3 (12.0) | 7 (19.4) |  |
| no | 11 (44.0) | 19 (52.8) |  |
| unknown | 11 (44.0) | 10 (27.8) |  |
| **Lethargic** |  |  | 0.533 |
| yes | 14 (56.0) | 24 (66.7) |  |
| no | 0 (0.0) | 2 (5.5) |  |
| unknown | 11 (44.0) | 10 (27.8) |  |
| **Rearing environment** |  |  | 0.194 |
| indoor | 5 (20.0) | 2 (5.6) |  |
| outdoor | 4 (16.0) | 9 (25.0) |  |
| free roaming | 15 (60.0) | 23 (63.9) |  |
| unknown | 1 (4.0) | 2 (5.6) |  |

The contingency table displays the characteristics of SFTSV-positive cats according to whether they survived or died. N (%): number of cases (percentage).
SFTSV, severe fever with thrombocytopenia syndrome virus. Statistical significance: \* $p < 0.05$ , \*\* $p < 0.01$ , \*\*\* $p < 0.001$ .

We compared the laboratory parameters between the surviving and dead groups separately in the SFTSV-positive and SFTSV-negative groups. (Figure 2, Tables S3 and S4). In the SFTSV-negative group, body temperature and the RBC of the fatal group were significantly lower than those in the surviving group (body temperature: p = 0.009; RBC: p = 0.027), and creatine phosphokinase (CPK) levels of the fatal group were significantly higher than those in the surviving group (p = 0.031). In the SFTSV-positive group, alanine aminotransferase (ALT) and AST levels in the dead group were significantly higher than those in the surviving group (ALT: p = 0.021; AST: p = 0.008). Regardless of the outcome, the SFTS- positive cases had severe leukopenia.

**Figure 2.**
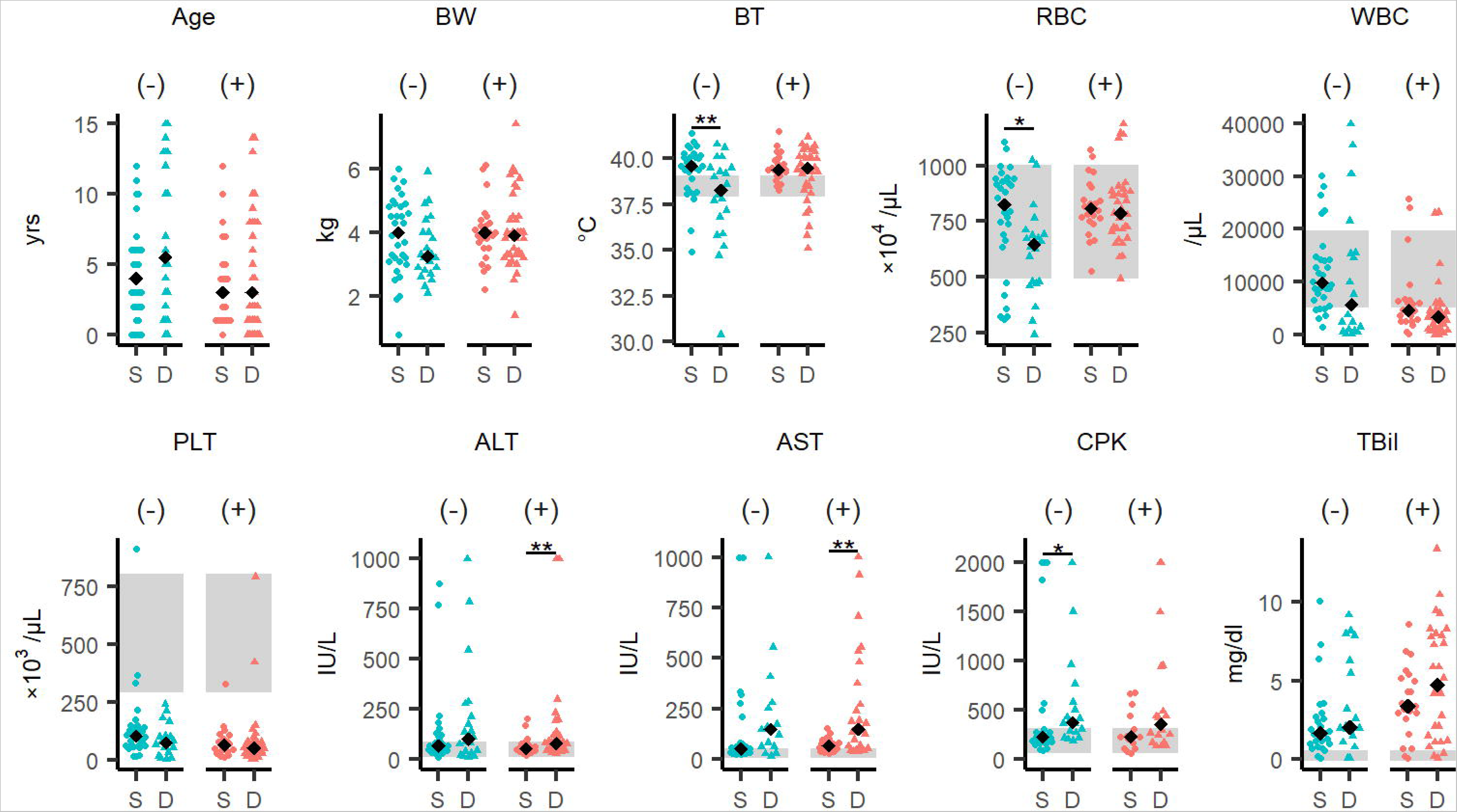
Differences in clinical characteristics between surviving and dead SFTSV-negative and SFTSV-positive cases. Comparison of individual data and laboratory parameters between the surviving and dead SFTSV-negative and SFTSV-positive groups. The black diamonds represent median values in each group. The blue circles or triangles represent the SFTSV-negative group, and the red circles or triangles represent the SFTSV- positive group. The circles and triangles represent individual values of the surviving group and dead group, respectively. The gray areas represent reference ranges for body temperature, RBC, WBC, PLT, ALT, AST, CPK, and TBil. “(–)” and “(+)” stand for “SFTSV-negative” and “SFTSV-positive,” respectively. Each variable was compared using the Wilcoxon rank-sum test, and statistical significance levels were corrected using the false-discovery rate (FDR). The level of statistical significance is shown: *p < 0.05, **p < 0.01, ***p < 0.001. See also Tables S3 and S4. Abbreviations: ALT, alanine aminotransferase; AST, aspartate aminotransferase; BT, body temperature; BW, body weight; CPK, creatine phosphokinase; D, died; PLT, platelets; RBC, red blood cells; S, survived; TBil, total bilirubin; WBC, white blood cell(s).

### Viral RNA levels of the SFTSV-positive cases

The amount of viral RNA in the SFTV-positive cases was measured using quantitative PCR (Figure 3). Serum-derived viral RNA levels were significantly higher in the fatal cases than in the surviving cases (p = 0.019). In contrast, there were no significant differences in the viral RNA levels extracted from eye swabs (p = 0.381), oral swabs (p = 0.525), or anal swabs (p = 0.510) between the two groups.

**Figure 3.**
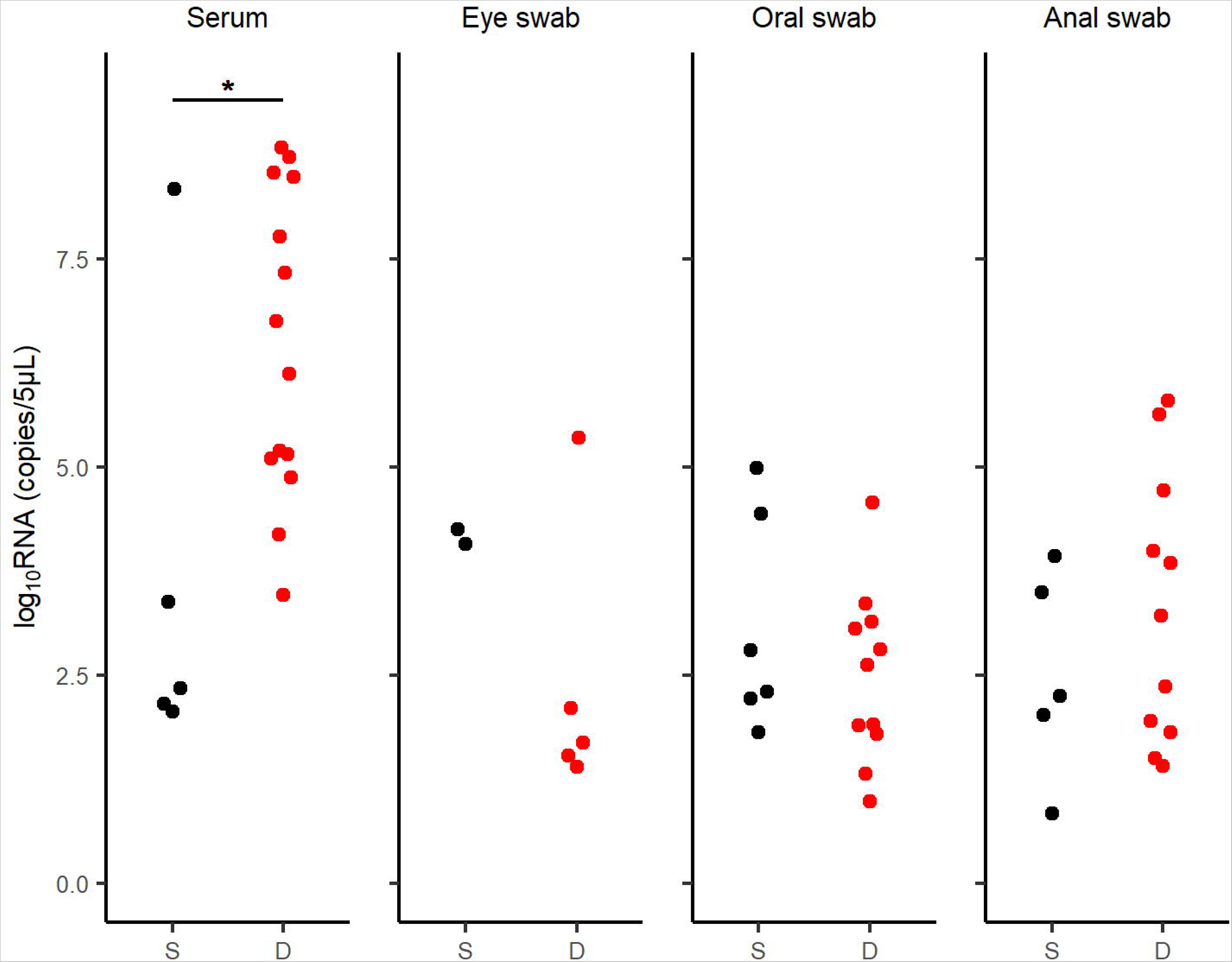
RNA levels in SFTSV-positive cases. RNA levels were detected in the serum, eye swabs, oral swabs, and anal swabs of SFTSV-positive cases. The black circles represent surviving cases and the red circles represent dead cases. The viral loads were compared using the Wilcoxon rank-sum test. (serum: p = 0.019, eye swab: p = 0.381, oral swab: p = 0.525, and anal swab: p = 0.510). Abbreviations: D, died; S, survived.

**Figure 4.**
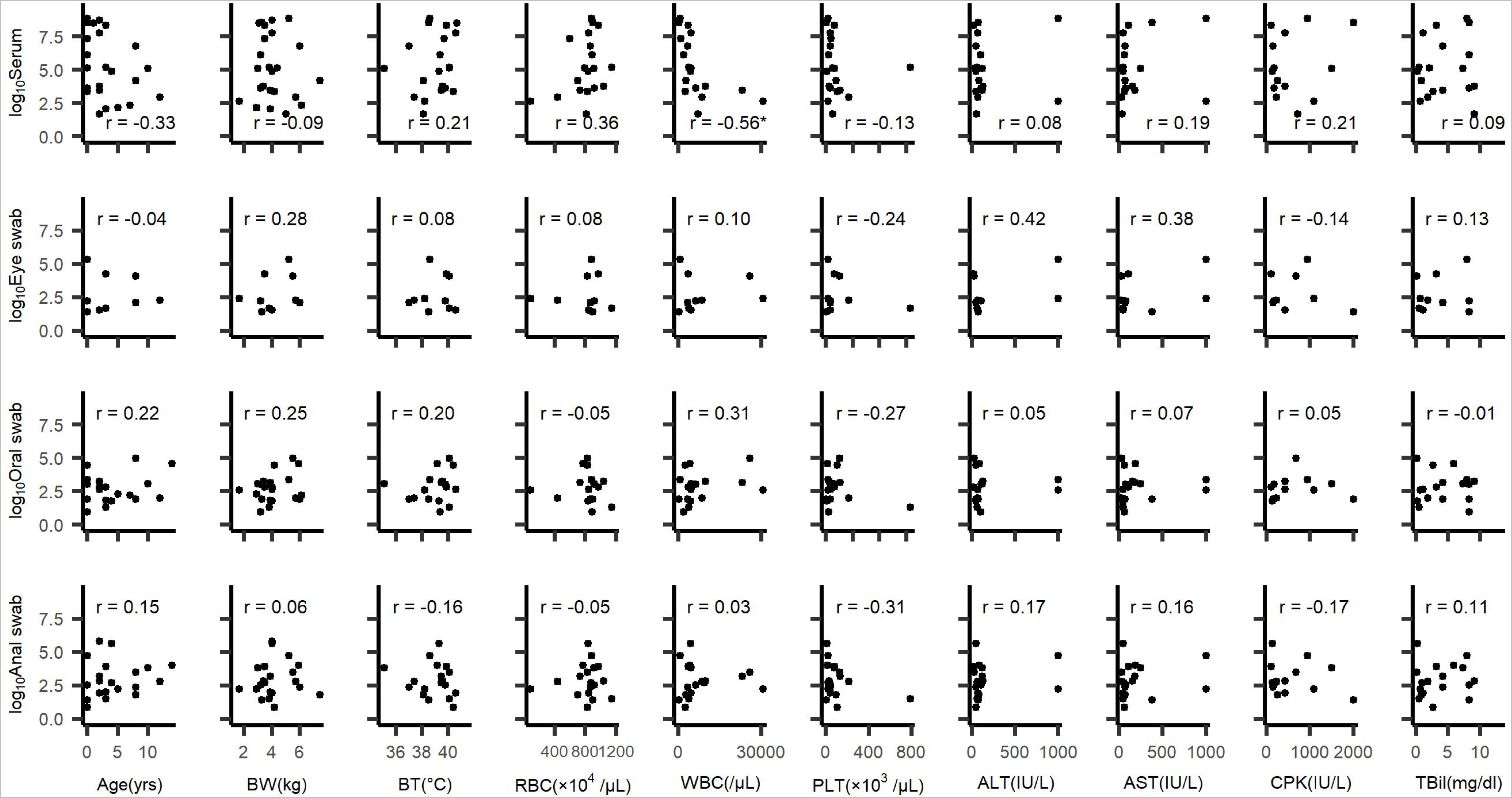
Correlations between RNA levels and epidemiological and clinical characteristics. Correlations between the viral load (detected in serum, eye swabs, oral swabs, and anal swabs) and variables such as age, BW, BT, RBC, WBC, PLT, ALT, AST, CPK, and TBil. The unit of RNA levels is copies/5-μL. The Pearson correlation (r) is shown at the upper left or lower right of each graph. Statistical significance: *p < 0.05, **p < 0.01, ***p < 0.001. Abbreviations: ALT, alanine aminotransferase; AST, aspartate aminotransferase; BT, body temperature; BW, body weight; CPK, creatine phosphokinase; PLT, platelets; RBC, red blood cells; TBil, total bilirubin; WBC, white blood cells.

### Antibody detection in SFTSV genome-negative serum specimens

SFTSV-reacting antibodies in the viral genome that were not detected in the 34 cat serums were evaluated by immunofluorescence analysis. The monoclonal antibody 4A10, used as a positive control, was able to detect specific staining for SFTSV at a dilution of 1:12800. In contrast, none of the RT-qPCR-negative cat serum samples showed any specific reaction against SFTSV at a dilution of 1:10 (Figure S1).

### Correlations between viral loads and clinical and laboratory parameters

The Pearson correlation was calculated between viral RNA loads and clinical or laboratory parameters in the SFTSV-positive groups. We found a negative correlation between serum viral load and WBC count (r = -0.56, p =0.013). Other parameters did not correlate with serum-derived viral RNA levels.

### Potential associations between SFTSV infection and lifestyle or clinical features in cases of feline SFTS

To examine the potential associations between SFTSV positivity and feline lifestyle or clinical features, we performed an analysis of associations (Figure 5). In the overall sample (n = 187), the SFTSV result (0: negative, 1: positive) was negatively associated with the WBC count and positively associated with the RBC count and TBil level. Age and tick parasitism (0: no bite; 1: bite) were negatively associated. Outcome (0: survived, 1: died) showed a negative association with the WBC count and a positive association with the AST level. In addition, body temperature positively correlated with ALT, AST, and CPK levels (Figure 5A).

**Figure 5.**
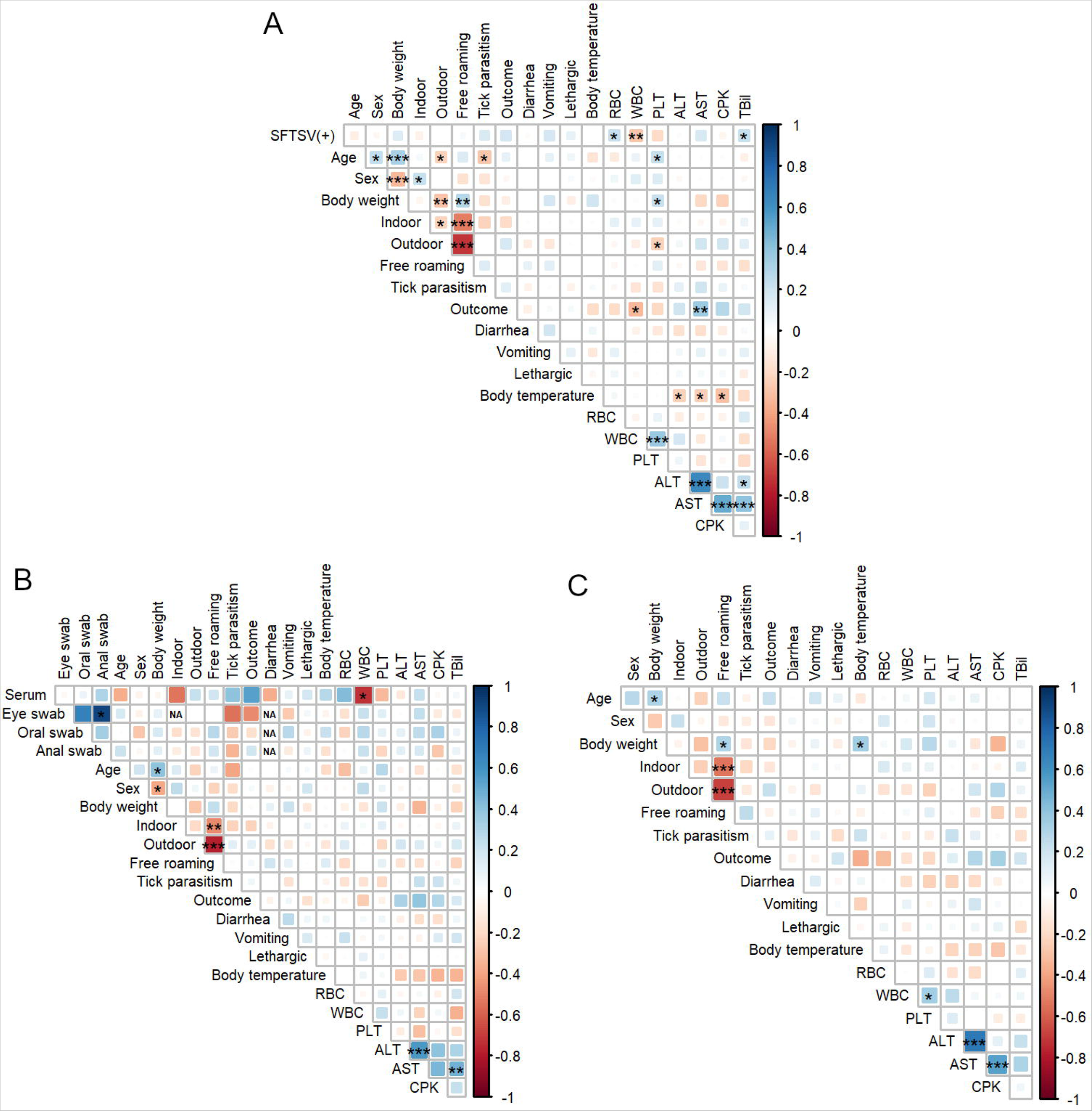
Spearman correlation matrixes in total cases, SFTSV-positive and SFTSV-negative cases. Spearman correlation matrixes of individual data, lifestyle, clinical data, and laboratory parameters in (A) total cases, (B) SFTSV-positive cases, and (C) SFTSV-negative cases. Results: SFTS positivity, sex (0: male; 1:female), indoor, outdoor, free roaming, tick parasitism, outcome, diarrhea, vomiting, and lethergy were set as dummy variables (0 vs. 1). Spearman correlation r-values are indicated using the square size and a heat scale. If the standard deviation is 0, the correlation coefficient cannot be calculated; therefore, the cell is labeled “NA.” The statistical significance levels, corrected using the false-discovery rate (FDR), are shown as squares; *p < 0.05, **p < 0.01, ***p < 0.001. Abbreviations: FDR, false-discovery rate; NA, not applicable; SFTS, severe fever with thrombocytopenia syndrome; SFTSV, severe fever with thrombocytopenia syndrome virus.

In the analysis of SFTSV-positive felines (n = 77), the WBC count was negatively correlated with serum viral RNA loads. Additionally, AST levels were positively correlated with ALT and TBil levels (Figure 5B).

In analysis of SFTSV-negative feline cases (n = 110), body weight showed a positive correlation with body temperature. Regarding other factors, there were positive correlations between WBC count and PLT, ALT and AST, and AST and CPK levels (Figure 5C).

### Scoring model for SFTS diagnosis

As mentioned above, SFTSV-positive cases demonstrated significantly higher levels of body weight, RBC, AST, and TBil and lower levels of WBC and PLT than SFTSV-negative cases (Figure 1). First, a receiver operating characteristic (ROC) curve analysis was performed (Table 3), and the optimal thresholds for each variable were set based on the cutoff values. Six variables were included in the stepwise multivariable logistic regression model to select more predictive variables (Table 4). The results showed that body weight ≥2.95 kg, RBC ≥697×10⁴ cells/μL, WBC <6370 cells/μL, AST ≥38 IU/L and TBil ≥4.05 mg/dL were independent factors for SFTSV positivity, but PLT <98×10³/μL was not. The Hosmer–Lemeshow test indicated that this logistic regression model fits well (p = 0.397).

**Table 3.** ROC curve analysis for independent factors.

| Parameter | AUC (95%CI) | Cutoff value | Sensitivity | Specificity | PPV | NPV |
| --- | --- | --- | --- | --- | --- | --- |
| **Body weight** | 0.586 (0.503-0.670) | 2.95 (kg) | 0.908 | 0.248 | 0.466 | 0.787 |
| **RBC** | 0.631 (0.546-0.716) | 697 ($\times 10^4/\mu\text{L}$) | 0.754 | 0.516 | 0.531 | 0.742 |
| **WBC** | 0.659 (0.576-0.742) | 6370 ($/\mu\text{L}$) | 0.732 | 0.602 | 0.571 | 0.756 |
| **PLT** | 0.615 (0.526-0.703) | 98 ($\times 10^3/\mu\text{L}$) | 0.788 | 0.441 | 0.500 | 0.745 |
| **AST** | 0.604 (0.511-0.696) | 38 (IU/L) | 0.918 | 0.354 | 0.514 | 0.852 |
| **TBil** | 0.632 (0.540-0.724) | 4.05 (mg/dL) | 0.523 | 0.775 | 0.630 | 0.690 |

ROC curve analysis was performed for five variables (Body weight, RBC, WBC, PLT, AST, and TBil) with $p < 0.05$ in the univariate analyses. AST, aspartate aminotransferase; AUC, area under the ROC curve; CI, confidence interval; NPV, negative predictive value; PLT, platelets; PPV, positive predictive value; RBC, red blood cells; TBil, total bilirubin; WBC, white blood cell(s).

**Table 4.** Multivariable logistic regression model for risk of SFTSV infection.

| Variable | SE | OR (95%CI) | P-value |
| --- | --- | --- | --- |
| Body weight ($\geq 2.95$ kg) | 0.750 | 3.11 (0.71-13.53) | 0.130 |
| RBC ($\geq 697 \times 10^4$ /μL) | 0.525 | 4.07 (1.73-13.50) | 0.004** |
| WBC ($< 6370$ /μL) | 0.486 | 4.07 (1.57-10.56) | 0.003** |
| AST ($\geq 38$ IU/L) | 0.655 | 3.24 (0.90-11.70) | 0.073 |
| TBil ($\geq 4.05$ mg/dL) | 0.497 | 2.40 (0.93-6.20) | 0.071 |

Five variables (Body weight, RBC, WBC, AST, and TBil) selected by the stepwise regression were entered into the multivariable logistic regression analysis. AST, aspartate aminotransferase; CI, confidence interval; OR, odds ratio; PLT, platelets; RBC, red blood cells; TBil, total bilirubin; WBC, white blood cell(s).
Statistical significance: \* $p < 0.05$ , \*\* $p < 0.01$ , \*\*\* $p < 0.001$ .

By integration of the five factors, namely body weight ≥2.95 kg, RBC ≥697×10⁴ cells/µL, WBC <6370 cells/µL, AST ≥38 IU/L and TBil ≥4.05 mg/dL, a scoring model was constructed with a possible range of 0 to 6 points (Figure 6A). Scores were assigned to each variable based on the β coefficients in the multivariable analysis. ROC curve analysis was performed for the scoring model, which indicated that the area under the curve (AUC) of this model was 0.828 (Figure 6B). This scoring model has a high sensitivity and may be a useful one for identifying SFTS infection based on clinical findings.

**Figure 6.**
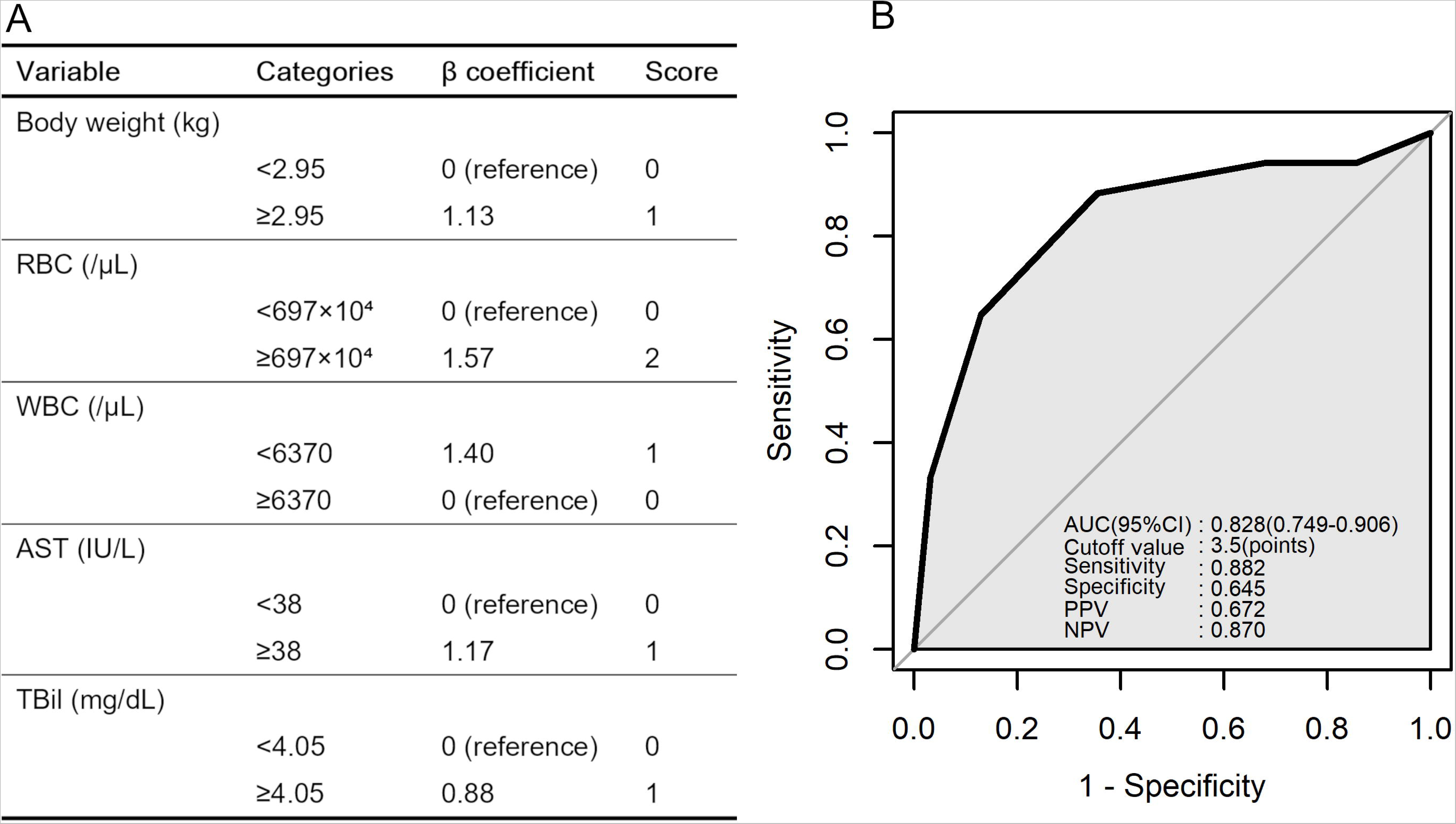
Scoring model for SFTS diagnosis. (A) Scoring model to predict infection of SFTSV in cats, based on the β coefficients in the multivariable logistic regression analysis. The scoring model has a possible range of 0 to 6 points. (B) ROC curve for the scoring model to predict SFTSV infection in feline cases. The results of the ROC curve analysis for the scoring model are shown on the lower right. Abbreviations: AUC, area under the curve; CI, confidence interval; NPV, negative predictive value; PPV, positive predictive value; ROC, receiver operating characteristic; SFTS severe fever with thrombocytopenia syndrome; SFTSV, severe fever with thrombocytopenia syndrome virus.

### Phylogenetic analysis of SFTSV viral genome sequence isolated from cats

Based on the diversity of the SFTSV M segment compared with the L and S segments,^3,33^ SFTSV M segment phylogenetic trees were constructed to understand the evolutionary relationship between the 16 newly isolated SFTSV strains and previous studies^23^ (Figure 7). The 15 SFTSV isolates (accession numbers: PP273701–PP273716) were classified as the J1 clade, whereas P2021-09 (accession number: PP273702) was classified as the J3 clade. Notably, our isolate, P2021-09, was closely related to the MN995270 isolates from a previous study, which were also classified as the J3 clade. Our analysis demonstrated no clear association between the outcome and viral genetic variation.

**Figure 7.**
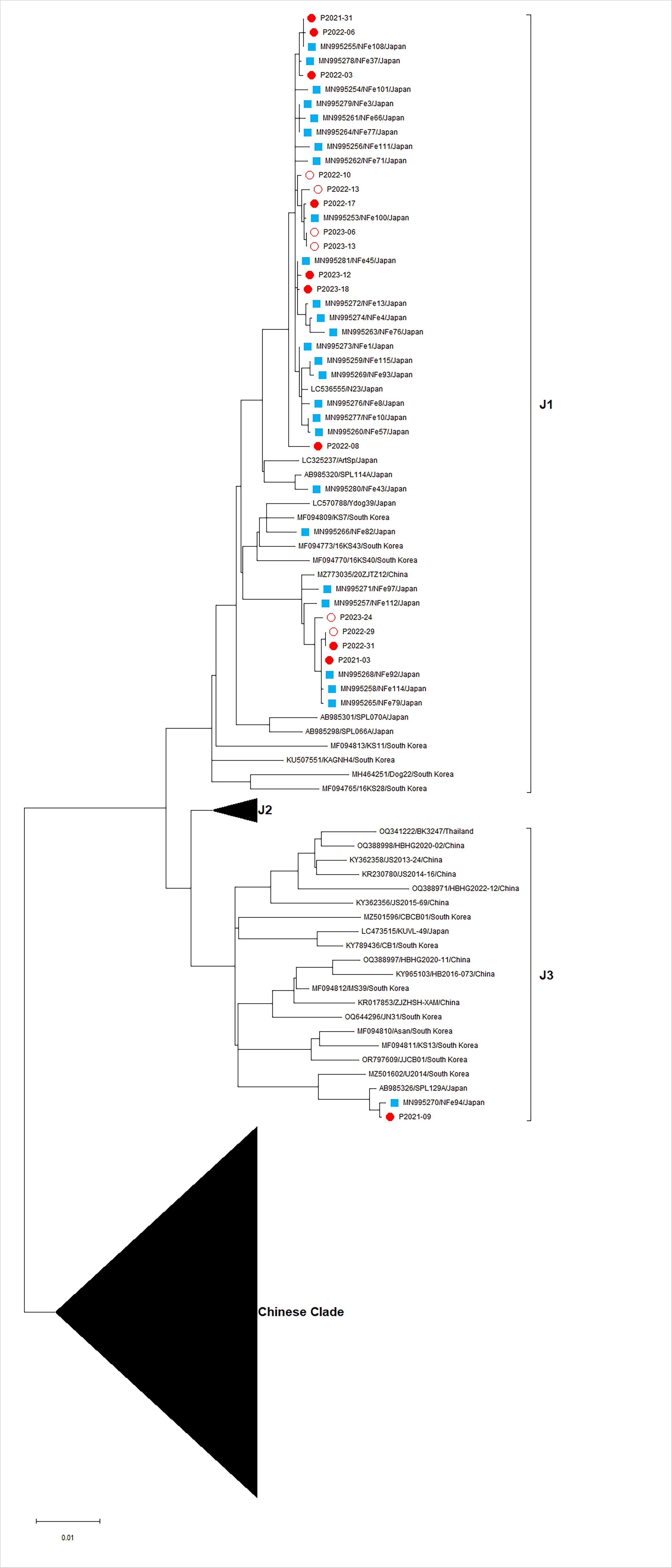
Phylogenetic tree analysis based on the SFTSV M segment. Maximum likelihood trees of nucleotide sequences from the M segment were constructed using MEGA11. The 16 sequences acquired in this investigation are depicted by circles, where white circles indicate the viruses isolated from survivors and red circles indicate those from fatal cases. The twenty-seven sequences obtained in our previous study are depicted by blue squares.

## Discussion

In this study, data from 187 cases of suspected feline SFTS were analyzed, and a comparison of SFTSV-positive and SFTSV-negative cases revealed that positive cases were characterized by heavier body weight, increased RBC counts, and AST and TBil levels, and decreased WBC and PLT counts. Furthermore, regarding the outcome of the SFTS-positive group, increased ALT, AST and higher serum RNA levels were identified in fatal cases.

Body weight was associated with SFTSV infection. Although obesity might be a potential risk for SFTSV infection, we need further investigation to clarify it because the body weight varies according to the cat breed, gender, and age. Although there was no significant difference of positivity ratio, vomiting was frequently observed in SFTSV infected feline cases, which has not been observed in humans. When bleeding tendency, which is a sign of SFTS, decreased RBC count was pronounced. The RBC count was slightly higher in SFTSV-positive feline cases than in SFTSV-negative cases; however, both were within the normal range. This might be due more to the effect of dehydration caused by fever and vomiting than bleeding. Notably, the WBC count decreased in SFTS-infected cases. Similar to human cases,^17^ cats also demonstrate leukopenia in SFTSV infection and are considered a useful differential marker for diagnosis. Decreased PLT counts, which is often observed in human SFTS cases,^17^ was pronounced in the SFTSV-positive feline cases. As we collected suspected SFTS cases, many of them had decreased PLT counts, even in SFTSV-negative cases. It is possible that some SFTSV-negative cases included other diseases with thrombocytopenia, such as feline immunodeficiency virus infection, hematologic disorders and tumors.^34^ In this regard, elevated levels of AST and TBil in SFTSV infection might be helpful for differential diagnosis. In the present study, we identified a significant difference in the TBil and AST levels, which is consistent with the results of previous studies.^11,35^ The liver damage might be induced by secondary pathological processes of SFTSV infection in cats, such as shock, hypercytokinemia, and hemophagocytosis.

For SFTS diagnosis, it is difficult to use RBC, WBC, TBil, PLT, AST, and TBil values alone because of the extremely small AUC values for the individual parameters in the ROC analysis. However, setting an optimal threshold for each and scoring them may provide a more efficient and practical threshold for SFTS diagnosis. We have tried several attempts and found optimal thresholds of over 3.5 points for SFTS diagnosis with 88.2% sensitivity and 64.5% specificity using Body weight ≥2.95kg, RBC ≥697×10⁴ cells/µL (2 points), WBC <6370 cells/µL (1 point), AST ≥38 IU/L (1 point), TBil ≥4.05 mg/dL (1 point). Our criteria for diagnosing SFTSV infection in cats need to be validated using clinical specimens.

In this study, ALT and AST levels were higher in the fatal cases than in the cats that survived. Elevated values of ALT,^36,37^ AST,^36–39^ and CK,^36–39^ have been reported in fatal human cases and these abnormalities might be indicators of disease severity and MOF. Serum viral RNA levels were higher in fatal cases than in surviving cases, as previously reported, and higher serum RNA levels were strongly associated with a fatal outcome.^11,40^ However, age was not positively correlated with the serum RNA load. This is consistent with the finding that age is not related to mortality in cats with SFTSV infection, in contrast with human cases.^36,41,42^ Therefore, age may not be a risk factor for SFTSV infection or death in cats. Overall, this study demonstrated that ALT, AST and serum viral RNA levels could be indicators of disease severity in cats with SFTSV infection. The timing of the nadir of the WBC count and the peak viral load were almost identical,^43^ and the finding of a negative correlation between the WBC counts and RNA levels is consistent with this finding.

In humans with SFTSV infection, an increase of IL-6, IL-8, IL-10, G-CSF and IFN-α are associated with an increased risk of death and are associated with viral load.^44^ In cats with SFTSV infection, high susceptibility to viruses may lead to amplification of the viral load and, consequently, to an exaggerated immune response elicited by SFTSV infection. Neither this study nor previous studies have been able to explain the severity of the disease based on differences in viral genetic diversity in cats with SFTSV infection.^23,45^ Notably, a recent study of human SFTS discovered one specific viral clade related to a higher case fatality rate.^46^ Thus, it is important to continue monitoring the relationship between viral genetic diversity and disease severity in feline cases.

## Limitations of the study

This study has some limitations. It did not collect repeated specimens over time, so was unable to characterize disease progression. Further studies are needed to obtain more accurate insights into disease progression and prognosis. In this study, we did not have a validation dataset to test the accuracy of our proposed scoring model; thus validation is required.

## Author contributions

All authors contributed to the intellectual content of this manuscript and approved the final manuscript as submitted. Conceptualization, Y.T; sample collection, Q.X, T.N, J.C.B, K.M.N, F.X.Y, S.I, D.H, M.M.N.T, K.M and Y.T; experiments, H.O, Q.X and Y.T; analysis, H.O, Q.X, T.N, and Y.T; funding acquisition, Y.T; drafted the initial manuscript, H.O, Q.X, and Y.T.

## Acknowledgments

The authors are grateful to Dr. Sho Miyamoto for his insightful comments, and Ayano Matsuzaki, Tomomi Kurashige, Megumi Tsubota, and Mika Ueda for their technical support. The authors are grateful to all cats and families who participated in this study and for their collaboration with veterinarians, animal hospitals (Sawamoto Inuneko Byouin, Tanigutchi Doubutsu Byouin, Shiroyama Doubutsu Byouin, Mine Doubutsu Byouin, Nagasaki Doubutsu Byouin, Kusunoki Doubutsu Byouin, Tamai Doubutsu Byouin, Hata Doubutsu Byouin, Fujii Doubutsu Byouin, Clover Doubutsu Byouin, Wada Doubutsu Byouin, Kaize Doubutsu Byouin, Taisuke Doubutsu Byouin, Yoshida Doubutsu Byouin, Misuna Juui Ika Iin, Pet no Byouin Katou, Isahaya Pet Clinic, Yoshioka Doubutsu Byouin, Shimabara Doubutsu Byouin, Momiji Doubutsu Byouin, Mori Doubutsu Byouin, Pet Clinic Iwakiri, Kaidu Doubutu Byouin, Hamada Doubutsu Byouin, Tsutsui Doubutsu Byouin, Tsuruno Doubutsu Byouin, Takekawa Inuneko Byouin, Tokoro Doubutus Byouin, Ariake Pet Clinic, Hinata Doubutsu Byouin, Azekari Doubutsu Byouin, Minami Inuneko Byouin, Urakawa Doubutsu Byouin, Higashinagasaki Pet Clinic, Parl Doubutsu Byouin, Taka Doubutsu Byouin, Yamamoto Inuneko Byouin, Iki Doubutsu Byouin, Matsumoto Juuika Iin, Nagasaki Cat Clinic, Hirota Inuneko Clinic, and Noa Doubutsu Byouin). The authors thank all the members of the Department of Virology, Institute of Tropical Medicine, Nagasaki University, and the funding partners listed below.

This research was supported by the Japan Agency of Medical Research and Development (AMED) under grant numbers JP23fm0208101, JP23fk0108656, JP23wm0125006, JP22wm0325023, JP22fm0208101, JP21fm0208101, JP21wm0325023, and JP20wm0323023; the Japan Society for Promotion of Sciences under grant numbers, 21K07059, 22KK0115, Takeda Science Foundation, MSD Life Science Foundation, the Naito Foundation, Kurozumi Medical Foundation, and Joint/Research Center on Tropical Disease, Institute of Tropical Medicine, Nagasaki University (2022-Ippan-12, 2023-Ippan-16).

## Availability of data and materials

The datasets generated and/or analyzed in the current study are available upon request to the responsible author.

## Supplemental information titles and legends

**Figure S1. Immunofluorescent analysis of SFTS-NP-mAb detection in viral RNA negative specimens.**

The SFTSV antigen slides were stained with SFTS-NP-mAb (4A10), which was used as a positive control, feline serum samples that were negative for viral RNA, and PBS as a negative control. The secondary antibodies used were Alexa Fluor 594 conjugated donkey anti-mouse and FITC-conjugated goat anti-feline. Nuclei were counterstained with Hoechst stain. Scale bar 20 μm. Partial results of the feline serum are displayed, and the mAb results are shown only for the 1:200, 1:800, 1:3200, and 1:12800 dilutions. Representative images are shown and the cat ID is indicated at the top. FITC, fluorescein isothiocyanate; PBS, phosphate-buffered saline; SFTS-NP-mAb, severe fever with thrombocytopenia syndrome virus monoclonal antibody; SFTSV, severe fever with thrombocytopenia syndrome virus.

**Table S1. Primers used in this study.**

Table S1 presents the comprehensive details of the primers used in this study. The table includes the following information for each primer: gene region, genome position, and nucleotide sequence of the primers and probes.

**Table S2. Comparison of the clinical characteristics of SFTSV-positive and SFTSV-negative cases, related to Figure1.**

Each variable was compared in the SFTSV-positive and SFTSV-negative cases using the Wilcoxon rank-sum test, and statistical significance is shown: *p < 0.05, **p < 0.01, ***p < 0.001. IQR, interquartile range; N, number of cases; Reference, reference range of each variable; SFTSV, severe fever with thrombocytopenia syndrome virus.

**Table S3. Comparison of the clinical characteristics between surviving and fatal SFTSV-negative feline cases, related to Figure 2**.

Each variable was compared in the surviving and fatal SFTSV-negative cases using the Wilcoxon rank-sum test, and the statistical significance is shown: *p < 0.05, **p < 0.01, ***p < 0.001. IQR, interquartile range; N, number of cases; SFTSV, severe fever with thrombocytopenia syndrome virus.

**Table S4. Comparison of the clinical characteristics between surviving and fatal SFTSV-positive feline cases, related to Figure 2**.

Each variable was compared in the surviving and fatal SFTSV-positive feline cases using Wilcoxon rank-sum test, and the statistical significance is shown;*p < 0.05, **p < 0.01, ***p < 0.001. IQR, interquartile range ; N, number of cases; SFTSV, severe fever with thrombocytopenia syndrome virus.

